# Antibiotics modulate activity of mouse and human dorsal root ganglia neurons

**DOI:** 10.64898/2026.09.10.750724

**Authors:** Ashley N. Plumb, Joseph B. Lesnak, Jane M. Brandon, Lucia M. Cardona, Mackenzie A. Simms, McKenna L. Pratt, Katie T. Ward, Kevin J. Tidgewell, Gregory Dussor, Theodore J. Price, Katelyn E. Sadler

## Abstract

Antibiotics are among the most prescribed medications worldwide, yet their direct effects on eukaryotic tissues remain largely unexplored. Here, we used *in vitro* calcium imaging to determine if clinically relevant concentrations of common antibiotics alter mouse and human dorsal root ganglion (DRG) neuron activity. We found that β-lactams, macrolides, and tetracyclines induce calcium flux in sensory neurons within minutes of application. Regardless of antibiotic class, neuronal responses depend on extracellular calcium entry. Pharmacological manipulations further revealed that cephalexin responses depend on TRPA1 activity whereas doxycycline responses require mitochondrial reactive oxygen species (ROS) production. These findings demonstrate that clinically relevant concentrations of antibiotics directly modulate sensory neuron activity through divergent mechanisms, thus expanding our current understanding of antibiotic side effects.

**Summary:** Clinically relevant antibiotics induce calcium flux in mouse and human sensory neurons through unique, drug-specific mechanisms.

## Introduction

Antibiotic use is both widespread and rapidly increasing, with global consumption estimated to exceed 70 billion annual daily doses by 2030 *(1)*. Antibiotics have transformed modern medicine and remain essential for treatment of bacterial infections, but their escalating use has contributed to a global antibiotic resistance crisis and increased recognition of adverse drug side effects including tendinopathy *(2)*, ototoxicity *(3)*, nephrotoxicity *(4)*, antibiotic-associated diarrhea *(5)* and visceral pain *(6)*. Although antibiotics are primarily classified by their bacterial mechanism of action, accumulating evidence indicates that these drugs also directly interact with host cells and influence eukaryotic physiology, independent of their antimicrobial activity. For instance, treatment with ciprofloxacin, a broad-spectrum fluoroquinolone, alters serum metabolites in germ-free mice, thus demonstrating that antibiotics directly manipulate host cell metabolism in the absence of a microbiome *(7)*. Somewhat unsurprisingly, several classes of antibiotics have also been shown to perturb the function of mitochondria *(8)*, eukaryotic organelles that are hypothesized to be evolutionarily descended from bacteria *(9)*. Given this, antibiotics may alter physiology in a broader range of host cells than previously appreciated.

Dorsal root ganglia (DRG) sensory neurons are uniquely positioned to detect and respond to systemic antibiotic exposure for several reasons. First, DRG neurons innervate all aspects of the gastrointestinal tract, where their peripheral terminals can detect changes in the local tissue environment following oral antibiotic administration. In fact, prior to absorption, luminal antibiotic concentrations exceed those detected in circulation, thereby exposing gut-innervating sensory neurons to high levels of these drugs. Second, because DRG lack a conventional blood-nerve barrier *(10)* and are therefore directly exposed to circulating drugs, systemic administration may modulate peripheral sensory neuron function in ways that contribute not only to acute gastrointestinal dysmotility and visceral pain, but also more persistent, potentially sub-clinical, body-wide symptomology. Finally, because of their incredibly long axons and associated high energy demands, DRG neurons contain large, tightly regulated populations of mitochondria that may be particularly susceptible to antibiotic perturbation *(11, 12)*. Thus, to determine if DRG neurons are antibiotic-sensitive, we exposed primary mouse (mDRG) and human (hDRG) DRG sensory neurons to four of the most frequently prescribed oral antibiotics – amoxicillin (β-lactam), azithromycin (macrolide), doxycycline (tetracycline), and cephalexin (β-lactam) – and measured acute effects on cellular signaling. Complementary *in vivo* observations were collected in mice to determine if drug-induced changes in cellular physiology manifested in observable behaviors.

## Results

### Clinically relevant concentrations of antibiotics directly activate mouse DRG (mDRG) neurons

To determine whether short-term exposure (**Fig. 1a**) to clinically relevant antibiotics alters sensory neuron function, we performed *in vitro* calcium imaging of mDRG neurons. Acute extracellular application of each antibiotic produced calcium flux within minutes (**Fig. 1b–e**). In contrast, vehicle-exposed neurons exhibited minimal activity, suggesting that responses were indeed the result of drug actions on mDRG soma. Antibiotic exposure increased intracellular calcium only in a subset of somata, therefore prompting us to quantify the proportion of responsive cells across multiple drug concentrations (**Fig. 1f-i**). Cephalexin induced calcium flux in small diameter (<27µm) mDRG neurons (*i.e,* those most likely to be nociceptors) in a concentration dependent manner (**Fig. 1f**), while medium and large diameter somata responses to cephalexin did not statistically differ from vehicle responses. Similarly, amoxicillin preferentially induced calcium flux in small-diameter neurons (**Fig. 1g**). Despite the 100-fold increase in drug concentration between the lowest and highest tested doses, approximately 17% of mDRG neurons exhibited responses to amoxicillin, regardless of concentration. Although the percentage of neurons that responded to azithromycin was lower than all other drugs tested, the highest concentration of azithromycin induced calcium flux in ∼10% of mDRG neurons, regardless of soma diameter (**Fig. 1h**). Finally, of all drugs tested, doxycycline produced the most robust, concentration-dependent responses in both large and small diameter neurons (**Fig. 1i**). Doxycycline responses were most notable in small diameter neurons; 45% of putative nociceptors exhibited calcium flux when exposed to 5 µg/mL doxycycline.

**Figure 1:**
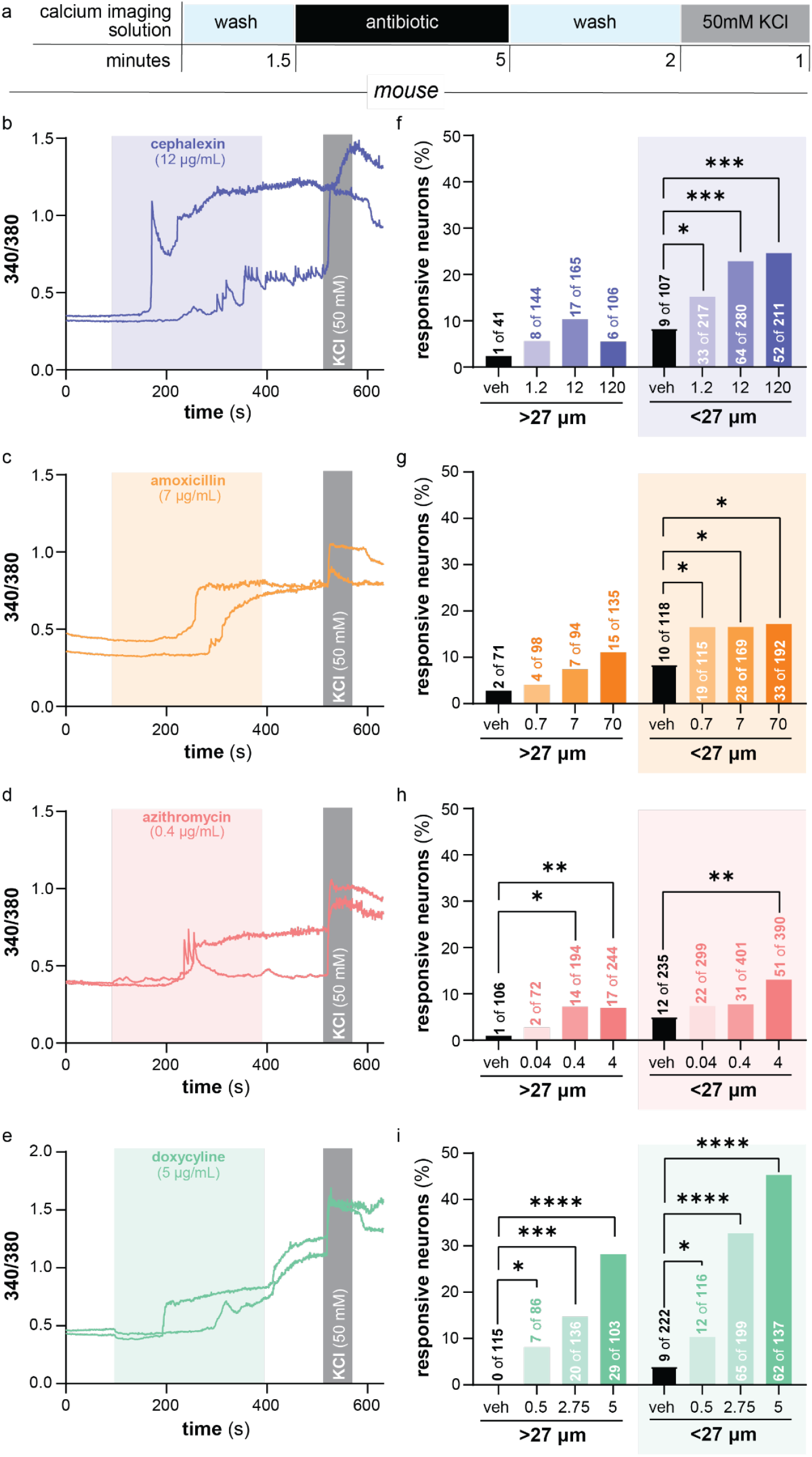
Commonly prescribed antibiotics directly activate mouse DRG neurons in vitro. **a.**, Timeline for calcium imaging experiments. Representative calcium transients from mDRG neurons exposed to **b.**, cephalexin (12 µg/mL), **c**., amoxicillin (7 µg/mL), **d.**, azithromycin (0.4 µg/mL), or **e.**, doxycycline (5 µg/mL). KCl (50 mM) was applied at the end of each recording to confirm neuronal viability. Quantification of neurons that responded to increasing concentrations of **f.**, cephalexin (1.2, 12, 120 µg/mL), **g.**, amoxicillin (0.7, 7, 70 µg/mL), **h.**, azithromycin (0.04, 0.4, 4 µg/mL), and **i.,** doxycycline (0.5, 2.75, 5 µg/mL). Large (>27 µm) and small diameter soma (≤27 µm) were analyzed independently. Numbers on top of or within bars indicate responsive neurons/total neurons analyzed; n=4 mice per drug. Statistical comparisons versus vehicle were performed using two-tailed Fisher’s exact tests within each size group; \**P*< 0.05, \*\**P* < 0.01, \*\*\**P* < 0.001, **** *P*< 0.0001.

Drug exposure only affected response frequency; in other words, antibiotic response magnitude and response kinetics did not differ across drug concentrations (**Fig. S1**). Because of the sustained increase in intracellular calcium in select cells, we performed a viable cell count following 5 minutes of exposure to the highest concentration of each drug to determine if antibiotic exposure induced excitotoxicity-associated cell death. Acute cephalexin and amoxicillin exposure induced modest cell death (**Fig. S2**). However, the percentage of non-viable cells in these assays was still lower than the percentage of cells that exhibited calcium flux upon drug exposure. In contrast, doxycycline and azithromycin had no effect on cell viability. Thus, in summary, antibiotics – regardless of drug class – induced calcium flux in putative mouse nociceptors.

### Antibiotic consumption induces acute changes in mouse pain-like behaviors and gastrointestinal motility

Because antibiotics induced calcium flux in a subset of mouse nociceptors, we next asked if oral consumption of clinically relevant doses of these drugs acutely altered rodent pain-like behaviors. Specifically, ongoing pain and abdominal mechanical sensitivity were measured in C57BL/6 mice following oral antibiotic administration using mouse grimace assessments and abdominal von Frey tests respectively (**Fig. 2a**). Regardless of drug class, antibiotic treatment induced facial grimacing 30 minutes following ingestion (**Fig. 2b-e**). Within each drug class, however, significant response variability was observed between animals. For example, some animals treated with low doses of doxycycline and cephalexin exhibited higher grimace scores than mice treated with higher drug doses. Summed responses over the first two hours following treatment revealed modest grimace score increases for all antibiotics (**Fig. 2f**).

**Figure 2:**
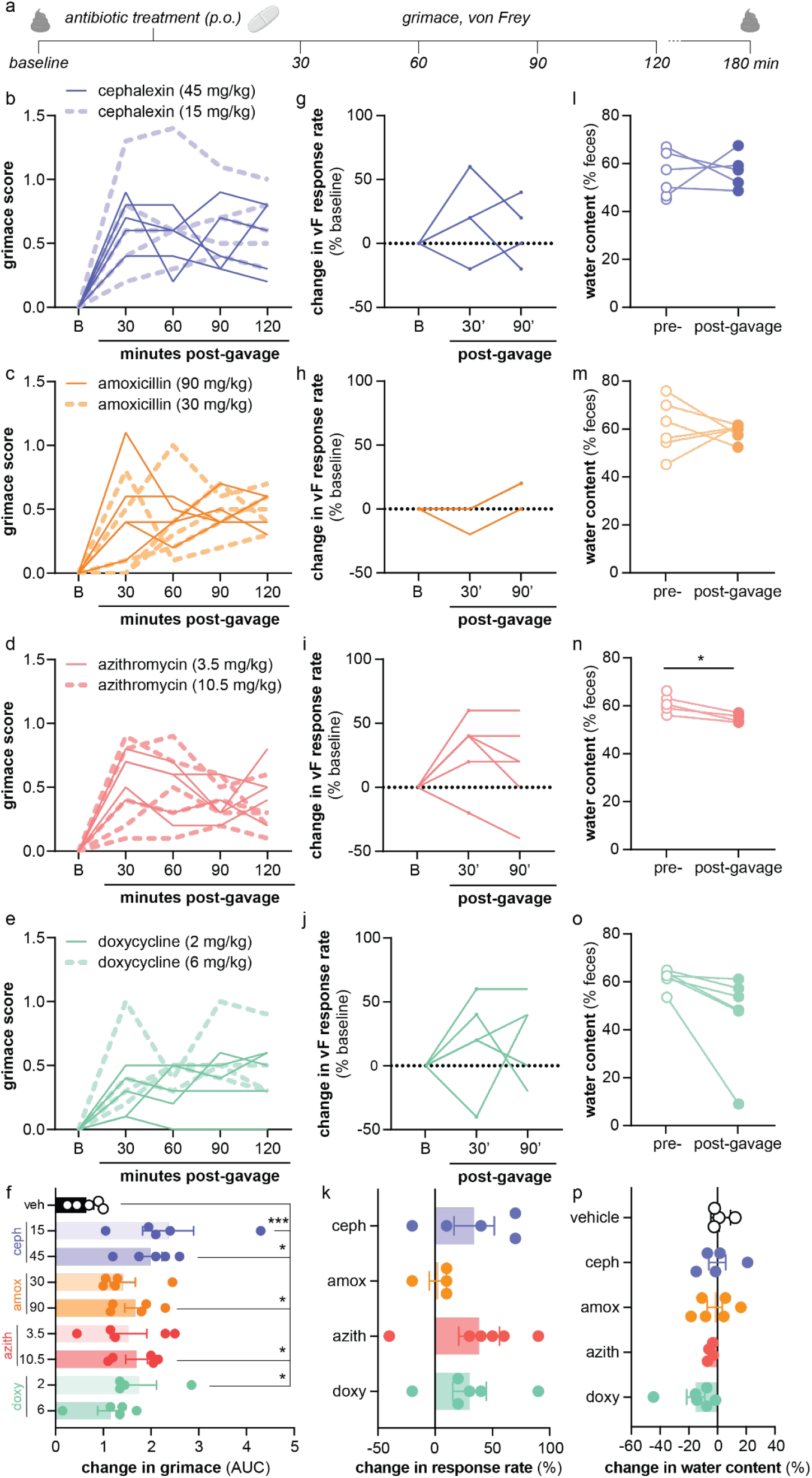
Oral antibiotic use induces modest increases in mouse pain-like behaviors. **a.**, Timeline for behavior experiments. Individual mouse grimace scores following oral administration of **b.**, cephalexin (15 or 45 mg/kg), **c.**, amoxicillin (30 or 90 mg/kg), **d.**, azithromycin (3.5 or 10.5 mg/kg), or **e.**, doxycycline (2 or 6 mg/kg). **f.**, Area under the curve (AUC) analysis of grimace assessments. Individual mouse changes in abdominal von Frey responsiveness following oral administration of **g.**, cephalexin (15 mg/kg), **h.**, amoxicillin (30 mg/kg), **i.**, azithromycin (2 mg/kg), or **j.**, doxycycline (3.5 mg/kg). **k.**, AUC analysis of von Frey assessments. Intra-animal variability in fecal water content prior to and 120-180 min following oral administration of **l.**, cephalexin (15 mg/kg), **m.**, amoxicillin (30 mg/kg), **n.**, azithromycin (3.5 mg/kg), or **o.**, doxycycline (2 mg/kg). **p**., Change in fecal water content following antibiotic treatment expressed as percent of baseline. Grimace AUC data analyzed separately for each antibiotic using one-way ANOVA followed by Dunnett’s multiple-comparisons test. Von Frey AUC data analyzed using two-tailed one-sample t tests against a theoretical mean of zero. Fecal water content analyzed using two-tailed paired t tests; *P < 0.05, **P < 0.01, ***P < 0.001, ****P < 0.0001.

In addition to non-evoked pain, we also assessed abdominal mechanical hypersensitivity for 90 minutes following antibiotic consumption. Similar to grimace, changes in abdominal sensitivity were most apparent 30 minutes following drug ingestion but varied substantially between animals and antibiotics (**Fig. 2g-j**). Unlike grimace scores which uniformly increased following administration of all antibiotics, abdominal hypersensitivity was not observed with all drugs; amoxicillin had essentially no effect on mechanical responsiveness (**Fig. 2k**). Finally, because gastrointestinal dysmotility is another prevalent side effect of antibiotic use, we assessed fecal water content as an indirect measure of diarrhea and/or constipation following antibiotic treatment. Of all antibiotics tested, azithromycin treatment was the only antibiotic that altered fecal water content post-treatment; stool collected shortly after azithromycin treatment contained less water suggesting that the drug induced a constipation-like phenotype (**Fig. 2l-p**). Thus, in summary, oral administration of clinically relevant doses of antibiotics induces modest increases in ongoing pain-like behaviors and abdominal hypersensitivity in mice that are likely not the result of altered gastrointestinal motility.

### Clinically relevant concentrations of antibiotics directly activate human DRG (hDRG) neurons

To complement mDRG imaging experiments, we next investigated whether clinically relevant, free/unbound plasma concentrations of antibiotics also alter human sensory neuron signaling. Using an imaging protocol that was identical to that used on mDRG (**Fig. 3a**), we observed that acute extracellular application of each antibiotic induced calcium flux in a subset of hDRG neurons (**Fig. 3b-e**). Given that our primary cell culture protocol biases smaller diameter (<60 µm) hDRG neuron survival, all recorded antibiotic-induced activity was in putative human nociceptors *(13)*. Like mDRG, a significant proportion of hDRG neurons exhibited calcium flux in response to acute exposure of amoxicillin, azithromycin, or doxycycline. When exposed to cephalexin, however, the proportion of responsive hDRG neurons did not differ from activation rates that were observed following vehicle exposure. Comparable proportions of hDRG and mDRG neurons responded when exposed to identical concentrations of amoxicillin or azithromycin. Doxycycline responses were blunted in hDRG with only 18% of hDRG neurons exhibiting responses during application of 5 µg/mL doxycycline, a concentration that induced responses in 45% of mDRG neurons (**Fig. 3f**). Finally, when compared to responses obtained during vehicle exposure alone, antibiotic responses did not differ in terms of magnitude (**Fig. 3g, 3h**) or latency (**Fig. 3i**). Together, these findings demonstrate that clinically relevant antibiotic exposure increases human nociceptor activity. These results highlight a previously unrecognized site of action for these widely used therapeutics.

**Figure 3:**
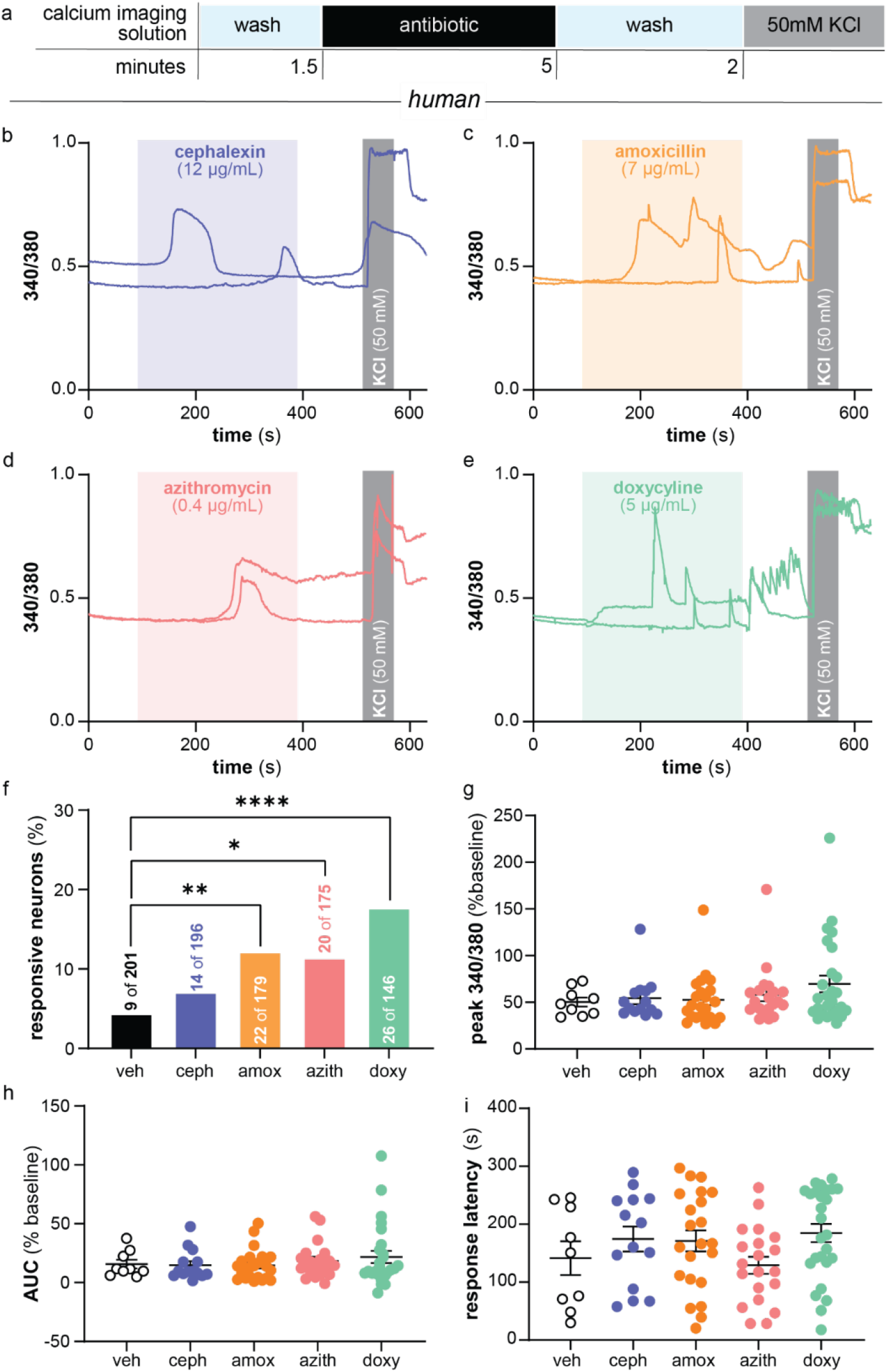
Commonly prescribed antibiotics directly activate human DRG neurons in vitro. **a.,** Timeline for calcium imaging experiments. Representative calcium transients from hDRG neurons exposed to **b.**, cephalexin (12 µg/mL), **c**., amoxicillin (7 µg/mL), **d.**, azithromycin (0.4 µg/mL), or **e.**, doxycycline (5 µg/mL). KCl (50 mM) was applied at the end of each recording to confirm neuronal viability. **f.**, Quantification of neurons that responded to each antibiotic. Comparison of antibiotic response **g.**, peak magnitude, **h.**, area under the curve (AUC), and **i.**, response latency between drugs. Responsivity analyzed via two-tailed Fisher’s exact tests; response magnitude, AUC, and latency analyzed using one-way ANOVA followed by Dunnett’s multiple-comparisons test. \**P*< 0.05, \*\**P* < 0.01, \*\*\**P* < 0.001, **** *P*< 0.0001.

### The molecular basis of antibiotic-induced neuronal activity varies between drug classes

We next sought to determine if different classes of antibiotics induce neuronal calcium flux through conserved or divergent molecular mechanisms. To begin, we measured antibiotic-induced neuronal activity in calcium-free conditions to assess the necessity of extracellular calcium influx. For these experiments, antibiotics were prepared in calcium-free buffer supplemented with 20 µM EGTA. Antibiotic solution was replaced with calcium-containing buffer immediately following drug exposure (**Fig. 4a**). As evidenced by representative traces, removal of extracellular calcium prevented almost all antibiotic-induced responses in mDRG neurons, regardless of the drug tested (**Fig. 4b-e**). However, upon re-addition of calcium to the extracellular milieu, many mDRG neurons exhibited quick, robust increases in calcium flux; note that these responses were not observed in the same proportion of vehicle-exposed neurons that were exposed to calcium-free buffer for the same period of time (**Fig. S3**). Quantification revealed that extracellular calcium was required for responses to all antibiotics tested (cephalexin, amoxicillin, azithromycin, and doxycycline; **Fig. 4f-i**). Upon re-addition of extracellular calcium, similar proportions of doxycycline-exposed mDRG neurons exhibited calcium transients as compared to mDRG neurons exposed to doxycycline in calcium-containing buffer (**Fig. 4i**). Intriguingly, re-addition of calcium following exposure to calcium-free solutions of cephalexin, amoxicillin, or azithromycin resulted in significantly more responsive neurons than if neurons were exposed to antibiotics in calcium-containing media (**Fig. 4f-h**). Based on this observation, cephalexin, amoxicillin, and azithromycin may exert their activity on voltage-sensitive, calcium-permeable channels that reside in the plasma membrane.

**Figure 4:**
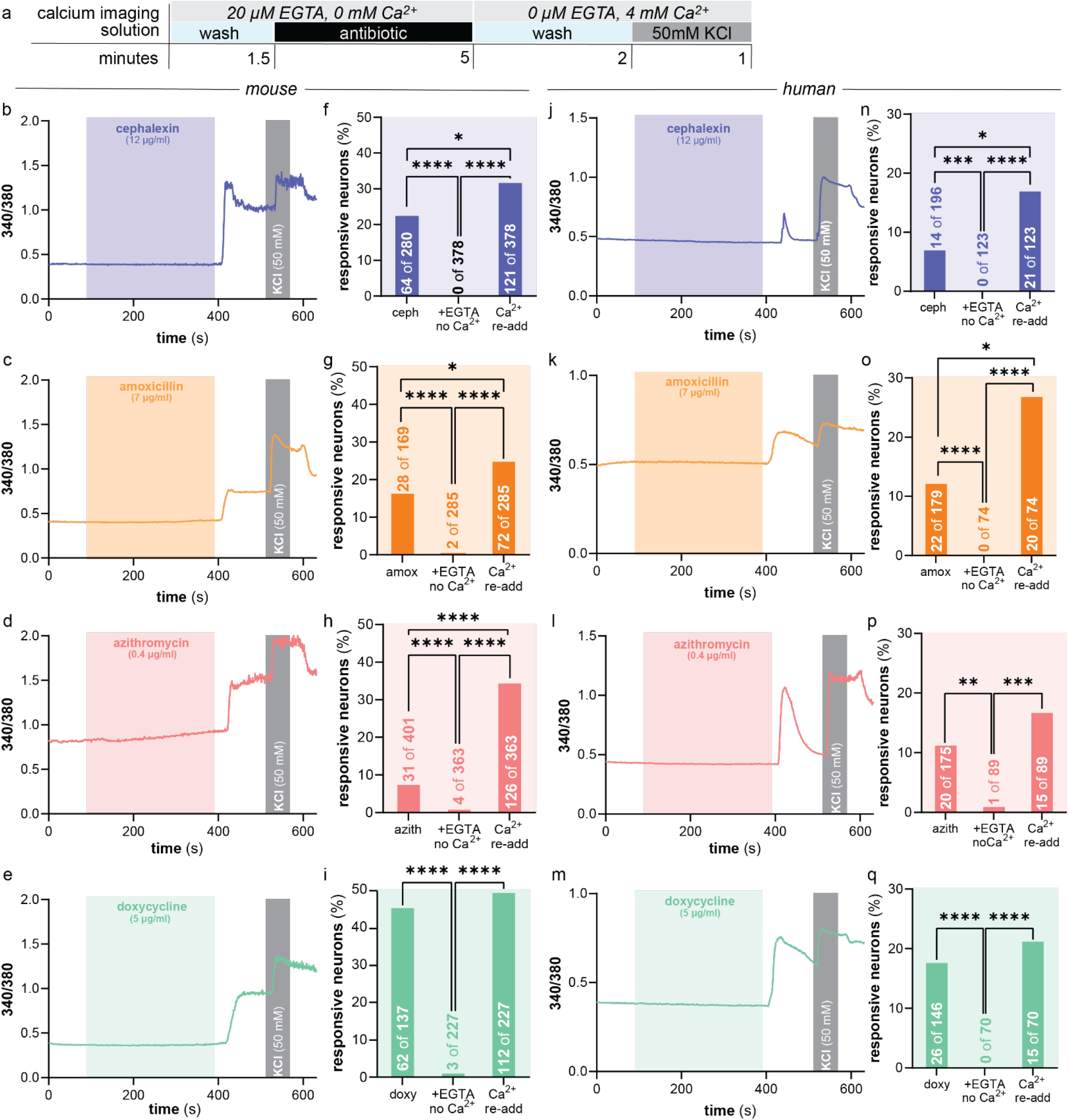
Antibiotic-associated activity in mDRG and hDRG neurons requires extracellular Ca^2+^ influx. **a.**, Timeline for calcium imaging experiments. Representative calcium transients from mDRG neurons exposed to **b.**, cephalexin (12 µg/mL), **c**., amoxicillin (7 µg/mL), **d.**, azithromycin (0.4 µg/mL), or **e.**, doxycycline (5 µg/mL) in calcium-free conditions (20 µM EGTA, 0 µM Ca^2+^) followed by re-addition of extracellular calcium. **f.-i.**, Quantification of mDRG neurons that responded during each phase of imaging (antibiotic alone data re-plotted from Figures 1 and 3, +EGTA/0 Ca^2+^, and Ca^2+^ re-addition). Representative calcium transients from hDRG neurons exposed to **j.**, cephalexin (12 µg/mL), **k**., amoxicillin (7 µg/mL), **l.**, azithromycin (0.4 µg/mL), or **m.**, doxycycline (5 µg/mL) in calcium-free conditions (20 µM EGTA, 0 µM Ca^2+^) followed by re-addition of extracellular calcium. **n.-q.**, Quantification of hDRG neurons that responded during each phase of imaging (antibiotic alone data re-plotted from figures 1 and 3, +EGTA/0 Ca^2+^, and Ca^2+^ re-addition). Statistical comparisons were performed using two-tailed Fisher’s exact tests; \**P*<0.05, \*\**P*<0.01, \*\*\**P*<0.001, \*\*\*\**P*<0.0001.

Calcium-free experiments were repeated with hDRG neurons to determine if antibiotic responses in these cells also depend on extracellular calcium influx. Like in mDRG neurons, essentially all antibiotic-associated activity in hDRG neurons was blocked when drugs were applied in calcium-free buffer containing 20 µM EGTA (**Fig. 4j-q**). Quantification of antibiotic responses again revealed that the percentage of cephalexin and amoxicillin sensitive neurons was increased when drugs were applied in a calcium-free solution followed by perfusion of calcium-containing media (**Fig. 4n, 4o**). Doxycycline responsivity of hDRG neurons was also similar to mDRG neurons in that it was not affected by extracellular solution calcium concentration (**Fig. 4q**). Interestingly, hDRG azithromycin sensitivity was notably different from mDRG. Application of calcium-free azithromycin solution activated the same percentage of hDRG neurons as calcium-containing azithromycin solution (**Fig. 4p**), but in mDRG, calcium-free azithromycin solution activated seven times as many neurons as calcium-containing azithromycin solution. Thus, hDRG neurons also require extracellular calcium for antibiotic-associated activity, but the precise molecular mechanisms underlying these responses may differ from those at play in mDRG neurons.

### Cephalexin and doxycycline activate mouse sensory neurons through distinct molecular mechanisms

Given the molecular clues generated by calcium-free imaging results, we next repeated antibiotic exposure in conjunction with application of ruthenium red, an inhibitor of many transient receptor potential (TRP) channels *(14)* and of mitochondrial calcium transport *(15)*(**Fig. 5a**). Interestingly, ruthenium red reduced the percentage of mDRG neurons that responded to cephalexin and doxycycline but had no effect on amoxicillin or azithromycin response frequency (**Fig. 5b-e**). In contrast, ruthenium red treatment decreased the percentage of hDRG neurons that responded to azithromycin, increased the percentage of hDRG neurons that responded to doxycycline application, and had no effect on the response frequency to amoxicillin or cephalexin (**Fig. 5f-i**). Integrating ruthenium red and calcium-free imaging results, the following hypotheses were generated regarding the molecular target of each antibiotic: (1) amoxicillin likely activates a calcium-permeable, voltage-sensitive, ruthenium red-insensitive channel in both mouse and human DRG; (2) azithromycin likely acts through different calcium-permeable channels in mouse and human DRG neurons; (3) cephalexin action in mDRG neurons may be partially mediated by a member of the TRP channel superfamily, and (4) doxycycline may alter mitochondrial activity in mDRG. Cephalexin and doxycycline hypotheses were subsequently tested with additional pharmacological manipulations on mDRG neurons.

**Figure 5:**
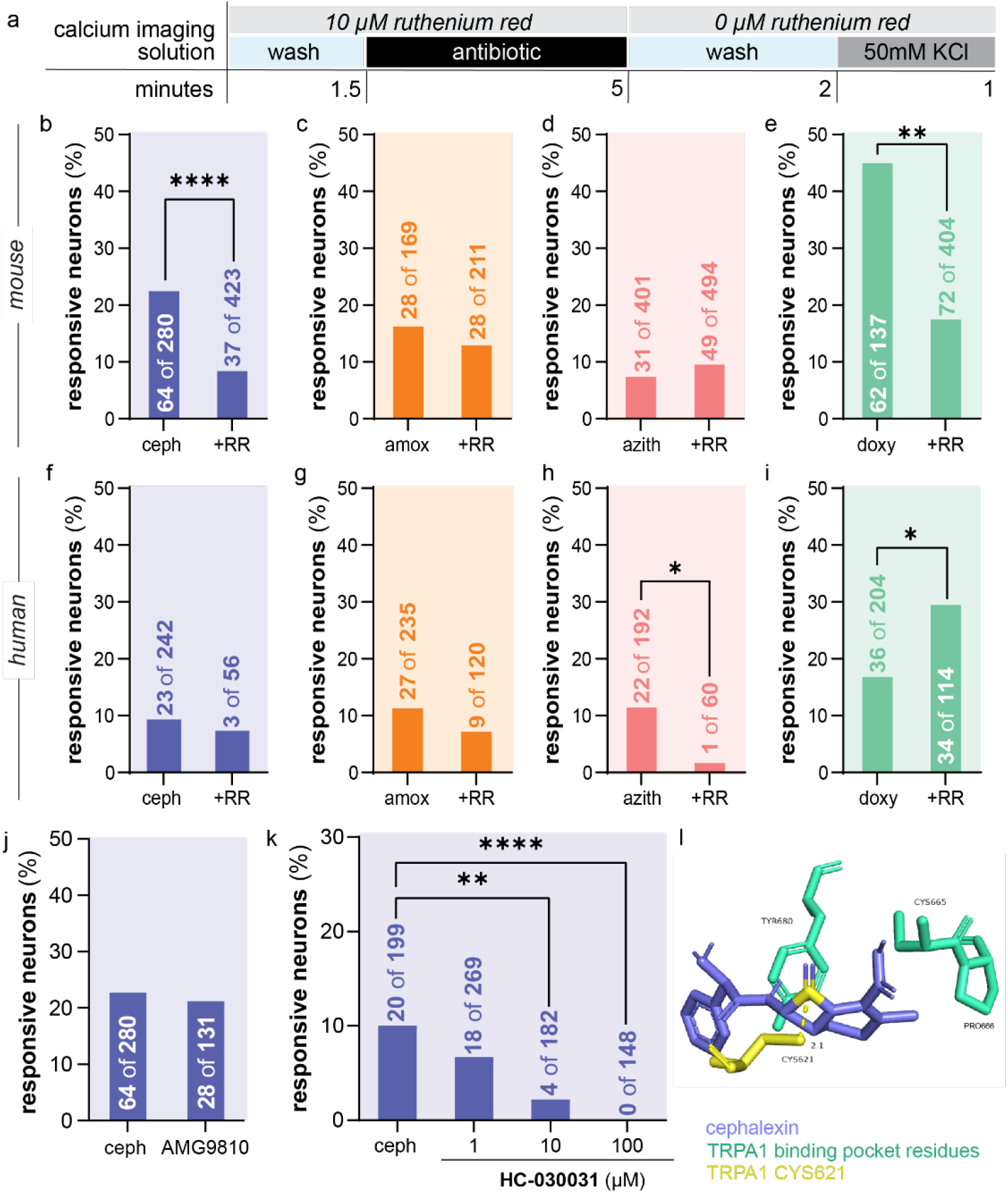
mDRG neurons respond to cephalexin via TRPA1 activity. **a.**, Timeline for calcium imaging experiments. Quantification of mDRG neurons that responded to **b.**, cephalexin (12 µg/mL), **c.**, amoxicillin (7 µg/mL), **d.**, azithromycin (0.4 µg/mL), or **e.,** doxycycline (5 µg/mL) when drugs were co-applied with ruthenium red (10 µM). Quantification of hDRG neurons that responded to **f.**, cephalexin (12 µg/mL), **g.**, amoxicillin (7 µg/mL), **h.**, azithromycin (0.4 µg/mL), or **i.,** doxycycline (5 µg/mL) when drugs were co-applied with ruthenium red (10 µM). Percentage of mDRG that responded to extracellular cephalexin exposure when co applied with **j.**, TRPV1 antagonist AMG9810 (10 µM) or **k.**, TRPA1 antagonist HC-030031 (1, 10, 100 µM). **l.,** Computational modeling of TRPA1 binding pocket interaction with cephalexin. Statistical comparisons were performed using two-tailed Fisher’s exact tests; \**P*<0.05, \*\**P*<0.01, \*\*\**P*<0.001, \*\*\*\**P*<0.0001.

Because cephalexin responses in mDRG were increased following exposure to calcium-free media and largely prevented with co-application of ruthenium red, we hypothesized that drug-related calcium flux may depend on activation of one or more TRP channels. After observing that co-application of the TRPV1 inhibitor AMG9810 had no effect on cephalexin-induced mDRG activity (**Fig. 5j**), we hypothesized that cephalexin may directly activate TRPA1 via covalent modification of channel cysteine residues *(16)*. Indeed, increasing concentrations of the TRPA1 antagonist HC-030031 dose-dependently reduced the percentage of mDRG neurons that responded to cephalexin (**Fig. 5k**). Computational modeling revealed that the β-lactam ring structure of cephalexin (*i.e.,* putative TRPA1 interaction site) adopts a suitable binding pose relative to TRPA1 CYS621 (**Fig. 5l**). Despite maintaining a similar β-lactam ring, no realistic poses exist for amoxicillin (**Fig. S4**). Before covalent binding, β-lactam drugs form a bent conformation, allowing nucleophilic attack to occur on the unhindered convex face of the ligand *(17, 18)*. Only one of amoxicillin’s predicted poses (**Fig. S4, Table S2**) adopted the correct position for binding but was not considered physically plausible due to steric clashes between CYS621 and the amine moiety. These docking data align with the inability of ruthenium red to block amoxicillin responses in either mDRG or hDRG neurons.

Like cephalexin, doxycycline responses in mDRG were similarly reduced following ruthenium red exposure but not impacted by removal of extracellular calcium. Therefore, we hypothesized that doxycycline may induce mDRG cytosolic calcium flux via mitochondrial activation and subsequent production of ROS. To test this hypothesis, mitochondrial ROS production was first measured in mDRG neurons using MitoSOX (**Fig. 6a**). Compared to vehicle treatment, doxycycline exposure increased both the proportion of MitoSOX-positive neurons (**Fig. 6b**) and the intensity of the MitoSOX signal in responsive neurons (**Fig. 6c**). As a positive control, co-incubation with the antioxidant N-acetylcysteine (NAC), significantly reduced both of these superoxide-dependent measures (**Fig. 6b, 6c**). Following this observation, we repeated doxycycline calcium imaging experiments with the addition of NAC in the extracellular solution. NAC incubation prevented a significant proportion of mDRG neurons from responding to doxycycline (**Fig. 6d**). In the few neurons that retained doxycycline sensitivity, time to reach the drug response threshold was significantly extended, further supporting the idea that ROS production is required for doxycycline-induced calcium responses (**Fig. 6e**). Collectively, these findings suggest that doxycycline-induced generation of mitochondrial ROS underlies mDRG neuron responses to this antibiotic.

**Figure 6:**
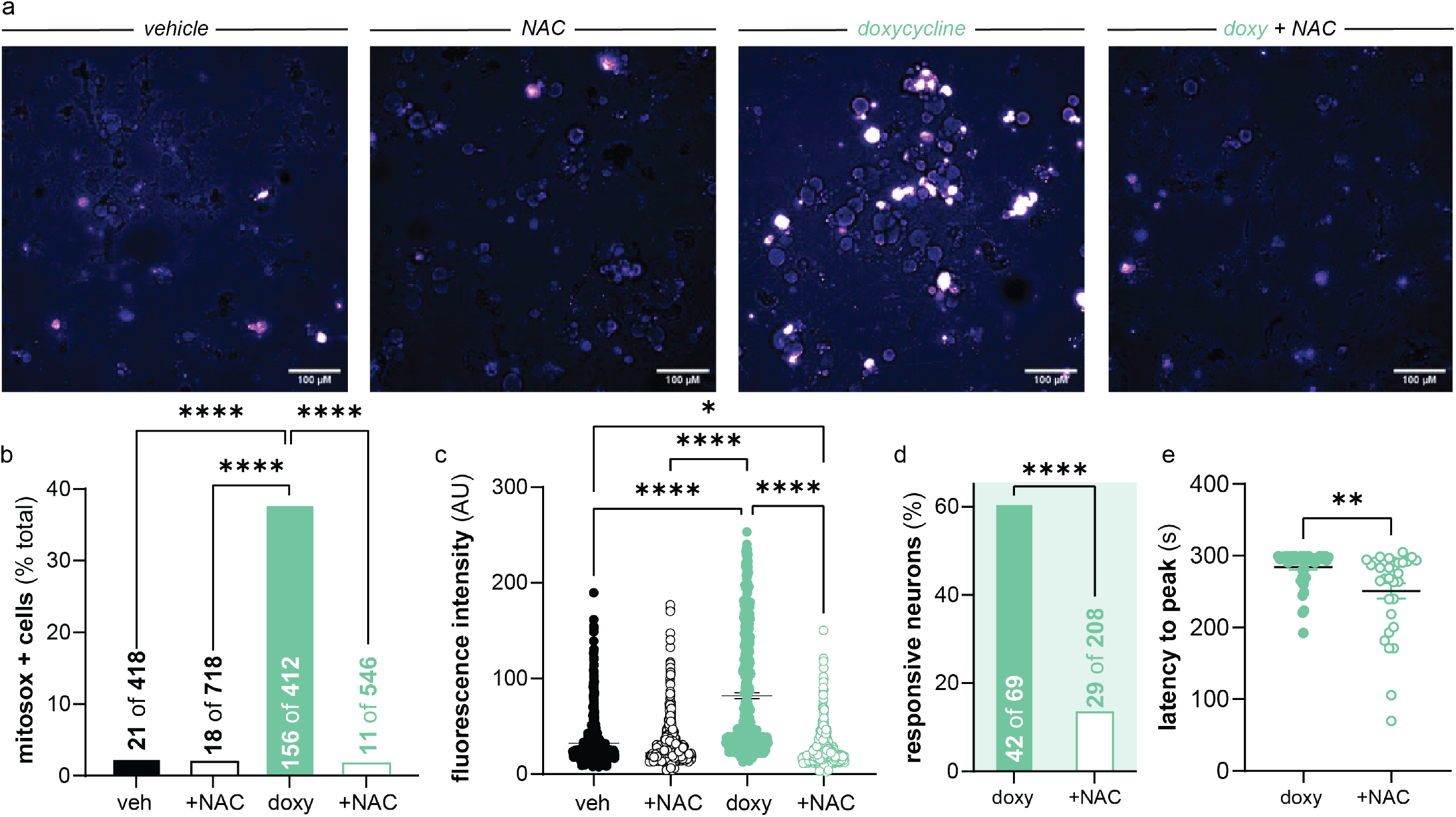
mDRG neurons mitochondrial ROS production required for doxycycline responses. **a.**, Representative images of MitoSOX signal (superoxide indicator) in mDRG neurons following exposure to vehicle or doxycycline (5 µg/mL) in the presence or absence of N-acetylcysteine (NAC; 10 mM). Quantification of **b.**, MitoSOX positive-cells and **c.**, MitoSOX signal intensity following exposure to vehicle or doxycycline in the presence or absence of N-acetylcysteine. **d.**, Quantification of mDRG neurons that responded to doxycycline when drug was applied in the presence or absence of N-acetylcysteine. **e.**, Time to reach doxycycline response threshold when drug is co-applied with or without N-acetylcysteine. Mitosox positive cells and doxycycline responsivity analyzed via two-tailed Fisher’s exact tests; latency analyzed using one-way ANOVA followed by Dunnett’s multiple-comparisons test. \**P*< 0.05, \*\**P* < 0.01, \*\*\**P* < 0.001, **** *P*< 0.0001.

## Discussion

Despite widespread use, the impact of antibiotics on eukaryotic host physiology – particularly the nervous system – remains incompletely understood. Here, we demonstrate that four of the most widely prescribed antibiotics directly modulate mDRG and hDRG neuron activity *in vitro*, independent of drug antimicrobial activity. Regardless of species, approximately 10-20% of small-diameter putative nociceptors exhibited calcium flux in response to extracellular application of antibiotic. The only exception to this observation was doxycycline which induced calcium flux in ∼45% of mDRG neurons. Mechanistically, all antibiotic responses depended on influx of extracellular calcium. Responsivity to all antibiotics was significantly reduced when drugs were applied in calcium-free conditions; rebound responses, which are presumably the result of continued drug-target engagement, emerged upon reintroduction of extracellular calcium. This indicates that antibiotic neuronal responses are not driven by intracellular calcium release alone but instead require influx from extracellular sources. For select drugs, response rates post-calcium re-addition exceeded rates observed when the same drug was applied in calcium-containing media. Critically, re-addition of extracellular calcium to vehicle-exposed neurons did not induce similar rates of neuronal activity. Collectively, these results support the idea that select antibiotics engage voltage-sensitive, calcium-permeable channels in primary sensory neurons.

Ruthenium red incubation further revealed that the exact molecular targets of each antibiotic vary between species and between drugs within the same class (*e.g.,* β-lactams cephalexin and amoxicillin). In mDRG neurons, cephalexin and doxycycline responses were significantly reduced by ruthenium red, but in hDRG neurons, ruthenium red only reduced azithromycin responsivity. Species differences, like the ruthenium red-sensitivity of azithromycin responses in hDRG, but not mDRG, may be explained by the species-specific channel pharmacology. For example, caffeine activates mouse TRPA1 channels and suppresses human TRPA1 activity. This difference arises from sequence variation within the N-terminus of the protein *(19, 20)*. Species differences may also be explained by differential TRP channel expression in mouse and human DRG neurons. *In situ* hybridization studies have identified key differences for several such superfamily members; *TRPV1 is* expressed in approximately twice as many hDRG neurons as mDRG neurons *(21), TRPA1* is expressed in almost five times as many mDRG neurons as hDRG neurons, and *TRPC5* is expressed in over five times as many hDRG neurons as mDRG neurons *(22)*. Given the relatively low expression of *TRPA1* in hDRG, it is thus unsurprising that ruthenium red exposure had no effect on cephalexin response rate in this species. Pharmacological manipulation of mDRG neurons, however, did support that TRPA1 – and not – TRPV1 is a primary target of cephalexin activity. TRPA1 can be activated via covalent bond formation between Cys621 and reactive functional groups on ligands like the aldehyde of cinnamaldehyde *(23)*, isothiocyanate of AITC *(24)*, or chloroacetyl of JT010 *(25)*. To explore whether the beta lactam ring of cephalexin can act as an electrophile and engage in a covalent bond with Cys621, we conducted docking studies of cephalexin, amoxicillin, and JT010 into human TRPA1 (PDB 6PQO). To mimic the covalent bond, we generated structures of the ligands attached to an end capped Cys before docking them into the crystal structure. Docking scores for all three were relatively similar (**Table S2**) however, cephalexin (2.8 A) and JT010 (3.4 A) are in closer proximity to Cys621 whereas amoxicillin (5.5 A) is potentially too far from the residue to form a covalent bond.

In addition to interacting with TRP channels, we also demonstrated that select antibiotics, namely doxycycline, alter neuronal calcium signaling through mitochondrial perturbation. Given the endosymbiotic origin of mitochondria, it is unsurprising that several antibiotic classes have been previously demonstrated to impair mitochondrial function *(26)*. Several classes of bactericidal antibiotics, including fluoroquinolones, aminoglycosides, and β-lactams, induce mitochondrial ROS production and associated oxidative stress in human mammary epithelial cells *(8)*. To our knowledge, the data presented here is the first report of a similar process occurring in primary sensory neurons. Sensory neurons have high metabolic activity *(27)*, and as such, mitochondrial perturbations can profoundly influence neuronal excitability. Specifically, we demonstrated that doxycycline induces mitochondrial ROS production and subsequent opening of calcium-permeable, plasma membrane localized channels in mDRG neurons. Given this initial intracellular site of action, doxycycline must pass through the plasma membrane to engage its target. The lipophilic nature of doxycycline permits passive membrane diffusion, but cephalexin – which presumably interacts with the cytosolic N-terminus of TRPA1 – likely depends on active transport mechanisms. Previous studies have identified PEPT1 and PEPT2 as cephalexin transporters *(28)*; the latter of these two proteins is expressed in hDRG *(29)*and mDRG *(30)* neurons, thus providing a candidate molecule for this process.

Our findings are consistent with – and an extension of – prior literature that documents the direct effects of antibiotics on eukaryotic cell physiology. Prior to this work, several antibiotic classes were associated with neurological side effects. For example, select fluoroquinolone and β-lactam drugs are neurotoxic due to their ability to inhibit GABA_A_ receptors *(31)*. Aminoglycosides interact with multiple targets in neurons; these drugs have been found to inhibit ASIC channel currents in primary sensory neurons and decrease central synaptic transmission by inhibiting voltage gated calcium channels *(32)*. Our work adds one additional ion channel and one intracellular organelle to the list of neuronal antibiotic targets, and suggests that antibiotics, as a collective drug class, have much broader effects on the mammalian nervous system than originally appreciated. Our results demonstrate that between 10-20% of hDRG neurons respond when exposed to clinically relevant concentrations of amoxicillin, azithromycin, or doxycycline. In learning this, one may then find themselves thinking back to the last time they took an antibiotic and re-examining whether there was a percept associated with this neuronal activity. Anecdotal accounts of antibiotic-associated pain and malaise are rampant, but here, for the first time, we demonstrate that antibiotic administration increased grimace scores – a correlate of multiple emotional states including pain *(33)*– in mice. An important caveat to these studies is the fact that antibiotics were administered to naïve animals that did not have an active bacterial infection. Although not always the case, antibiotics are most frequently prescribed to individuals with active infections that are often accompanied by – and in fact, frequently recognized by – localized pain. As such, it is possible that when ingested by patients, low level, and potentially widespread antibiotic-induced neuronal activity, either does not cross the perceptual threshold or is masked by the noxious sensations arising from the infection site. This “sub-threshold” neuronal activity should not be castoff, however. Long-term or repeated antibiotic use like that which occurs in sickle cell disease *(34)*, chronic Lyme disease *(35)*, and chronic pelvic pain disorders among other conditions *(36)*, may alter sensory neuron plasticity and ultimately contribute to the chronic pain experienced by those patients.

In addition to providing further support for better antibiotic stewardship, our findings have critical implications for the use of tetracycline derivatives in experimental systems. Tetracyclines, including doxycycline, are widely used in Tet-ON/Tet-OFF inducible gene expression systems. Here, we demonstrate that *in vitro* exposure to doxycycline induces calcium flux in up to 45% of mDRG neurons. This effect must be considered, especially when gene expression is being manipulated in rodent neurons. In addition, minocycline – another tetracycline that was originally developed as an antibiotic – is now widely used in preclinical research laboratories and clinical settings due to its microglial-mediated anti-inflammatory properties. Although not tested in the present experiments, it would be unsurprising if minocycline exposure also induced calcium flux in primary sensory neurons like its tetracycline relative. Lastly, because our data demonstrate that a wide range of antibiotics induce calcium flux in sensory neurons, potential effects of antibiotic exposure during sensory neuron culture protocols must be appreciated. Exposure to these drugs could alter basal neuronal physiology and subsequent responses to experimental stimuli.

Several limitations should be considered when evaluating these data. First, the majority of experiments herein were performed on dissociated neuron cultures that do not fully recapitulate the *in vivo* environment where antibiotics may exert additional activity on immune cells, glia, and the microbiota. Second, although ruthenium red sensitivity provides insight into potential channel involvement, it is a broad inhibitor, and future studies will be required to identify the specific ion channels mediating additional ruthenium red effects that were not explored in detail. Third, although our data demonstrate acute changes in neuronal function following antibiotic exposure, it is unclear if there are long-term consequences of antibiotic treatment on neuronal activity and plasticity. Future studies should continue to define the molecular players engaged by these and other antibiotic classes in order to minimize drug effects on eukaryotic physiology.

## Materials and Methods

### Animals

C57BL/6J mice were used for all experiments. Colony founders were purchased from The Jackson Laboratory then bred in house. Animals were maintained on a 12-hour light/dark schedule with *ad libitum* access to rodent chow and water. All animal experiments and procedures were performed in accordance with National Institutes of Health guidelines and were approved by the Institutional Animal Care and Use Committee at the University of Texas at Dallas (Richardson, TX; protocol no. 2022-0088).

### Primary sensory neuron isolation and culture

#### Mouse DRG neuron isolation and culture

Adult C57BL/6J mice (27 male and 5 female) were deeply anesthetized with isoflurane and euthanized by decapitation. DRG were rapidly dissected, then transferred to cold Hank’s Balanced Salt Solution (HBSS). Thoracolumbar (T10–L2) and mid-lumbar (L3–L4) DRG contain the cell bodies of afferents that innervate the small intestine and colon *(37)*. As such, bilateral T12-L2 and L3-L4 ganglia were collected and processed separately for calcium imaging experiments. Preliminary analyses revealed no differences in antibiotic responsiveness between thoracolumbar and lumbar neurons; therefore, data were combined for final analyses. Dissected ganglia were cultured as previously described *(22)*. Briefly, cells were incubated in DMEM containing collagenase type IV (final concentration of 1 mg/mL) for 40 minutes at 37°C. Following collagenase digestion, tissue was washed and incubated in DMEM containing trypsin (final concentration of 0.05%) for 45 minutes at 37°C. Enzymatic digestion was terminated by addition of heat-inactivated horse serum (0.5 mL) and ganglia were washed with DMEM prior to mechanical dissociation. Cells were gently triturated in complete media consisting of DMEM/F12 supplemented with 10% heat-inactivated horse serum, 2mM L-glutamine, and 1% glucose and plated onto laminin-coated glass coverslips. Note that antibiotics were not included in any step of the culture procedure. Cultures were maintained at 37°C in 5% CO2. After 1-2 hours of adherence, 1 mL of complete media was gently added to each well. Neurons were used for calcium imaging experiments within 24 hours of plating.

#### Human DRG Isolation and Culture

Human DRG were surgically recovered from organ donors in partnership with the Southwest Transplant Alliance. Tissue procurement procedures were approved by the University of Texas at Dallas Institutional Review Board under protocol Legacy-MR-15-237. Informed consent for research tissue donation from an organ donor’s first-person consent (driver’s license or legally binding document) or legal next of kin was obtained by the Southwest Transplant Alliance. The United Network for Organ Sharing approves all policies for screening and consent. The Southwest Transplant Alliance adheres to the standards and procedures established by the US Centers for Disease Control and these procedures are reviewed biannually by the Department of Health and Human Services. Circulation of any pertinent donor medical information strictly follows HIPAA regulations. Human DRG were surgically removed as previously described within 4 hours of cross-clamp *(38)*. DRG were immediately placed into either artificial cerebrospinal fluid (<1 hour) or Hibernate A supplemented with 1% N-2, 2% NeuroCult SM1, 1% penicillin/streptomycin, 1% Glutamax, 2mM Sodium Pyruvate, and 0.1% Bovine Serum Albumin (1-16hrs) at 4°C. Temporary storage of full DRG bulbs in Hibernate A solution prior to dissociation results in DRG neuronal cultures with comparable electrophysiology and calcium imaging data as freshly dissociated neuronal cultures *(39)*. A total of 38 DRGs were used from 14 organ donors (9 male and 5 female). Demographic information is provided in **Table S1**.

Human DRG cultures were prepared as previously described *(38)*. Briefly, the bulb of thoracic and lumbar DRG were exposed by trimming off excess connective tissue, fat, and nerve roots. The bulb of each DRG was diced into 3 × 3mm sections and placed in 5 mL of pre-warmed digestion enzyme consisting of 1 mg/mL of Stemxyme I, 0.1 mg/mL of DNase I, and 10 ng/mL of recombinant human β-NGF in HBSS without calcium and magnesium. The tubes were placed in a 37°C shaking water bath with light trituration every hour until DRG dissociated (∼4 hours). Samples were filtered through a 100 µM mesh strainer and the resultant cell suspension was gently layered over a 15 mL tube containing 3 mL of 10% BSA in HBSS. Tubes were centrifuged at 900g for 5 minutes at room temperature. The supernatant was aspirated and the pellet was resuspended in prewarmed BrainPhys® media containing 1% penicillin/streptomycin, 2% NeuroCult SM1, 1% Glutamax, 1% N-2, 10ng/mL β-NGF, 2% HyClone™ Fetal Bovine, and 0.1% 5-Fluoro-2′-deoxyuridine. Cells were plated on either 12mm coverslips or glass bottom plates pre-coated with 0.1mg/mL of poly-D-lysine depending on the experimental assay. Cells were incubated at 37°C and 5% CO2 for 3 hours to allow for adherence. Following adherence, wells were flooded with prewarmed media, and half media changes were performed every other day. Twenty-four hours prior to calcium imaging, DRG cultures were starved of antibiotics by completing a full media change to DRG media lacking penicillin/streptomycin.

### In vitro calcium imaging and pharmacological manipulations

Calcium imaging was performed on mouse and human DRG neurons using the ratiometric calcium indicator Fura-2 acetoxymethyl ester (Fura-2 AM). Neurons were loaded with Fura-2 AM (2.5 µg/mL for mouse; 3 µg/mL for human in 2% bovine serum albumin) in extracellular buffer for 45 minutes at room temperature followed by a 15-minute de-esterification period. Extracellular buffers were prepared for mouse and human cell imaging. Mouse buffer contained 150 mM NaCl, 5.6 mM KCl, 2 mM CaCl_2_, 1 mM MgCl_2_, 1 mM HEPES, and 8 mM glucose. Human buffer contained 145 mM NaCl, 3 mM KCl, 2.5 mM CaCl_2_, 1.2 mM MgCl_2_, 10 mM HEPES, and 7 mM glucose (pH 7.4 ± 0.03; 320 ± 3 mOsm). Cells were imaged using a Nikon Eclipse Ti200 inverted microscope. Fura-2 was alternately excited at 340 and 380 nm, and emission was collected at 510 nm. Images were acquired every 500 ms and intracellular calcium levels were expressed as a 340/380 nm fluorescence ratio. Regions of interest were drawn around neuronal soma and exported using NIS-elements software, and analysis was performed using a custom-designed script. Only neurons that responded to 50 mM KCl at the end of each experiment were included in analysis. A neuron was classified as antibiotic responsive if the peak 340/380 ratio increased by ≥25% above baseline.

#### Antibiotic application

Antibiotics were freshly prepared in extracellular buffer and applied via gravity-driven perfusion for 5 minutes. For mouse DRG experiments, three concentrations were tested for each antibiotic: amoxicillin (0.7, 7, 70 µg/mL), doxycycline (0.5, 2.75, 5 µg/mL), cephalexin (1.2, 12, 120 µg/mL), and azithromycin (0.04, 0.4, 4 µg/mL). For human DRG experiments, neurons were exposed to concentrations approximating clinically relevant peak free (unbound) serum levels: amoxicillin (7 µg/mL) *(40, 41)*, doxycycline (5 µg/mL) *(42, 43)*, cephalexin (12 µg/mL) *(44)*, and azithromycin (0.4 µg/mL) *(45)*. These concentrations were selected based on reported serum ranges and the pharmacologically active unbound fraction of each drug. Higher concentrations of doxycycline were not tested due to ultraviolet autofluorescence that interfered with Fura-2 imaging. To assess neuronal viability following antibiotic exposure, cultured DRG neurons were incubated with antibiotics at clinically relevant concentrations for 5 minutes. Cells were then stained with trypan blue (0.4%) and imaged using brightfield microscopy. The percentage of trypan blue–positive neurons was quantified relative to total neurons per field.

#### Pharmacological manipulations

To assess the contribution of extracellular calcium influx to antibiotic-induced neuronal activation, experiments were performed in calcium-free extracellular buffer in which CaCl_2_ was omitted and EGTA (20 µM final concentration) was included. Neurons were continuously perfused with this calcium-free, EGTA-containing solution during baseline recording and antibiotic exposure. To evaluate the involvement of TRP channel–mediated signaling pathways, Ruthenium Red (10 µM final concentration) was included in the extracellular bath solution during baseline perfusion and antibiotic application. EGTA and Ruthenium Red experiments were conducted on both mouse and human DRG neurons. To determine whether oxidative stress contributed to doxycycline-induced neuronal activation, N-acetylcysteine (NAC; 10 mM final concentration) was included in the extracellular bath solution during doxycycline exposure. To assess the involvement of TRPA1 in cephalexin-induced neuronal activation, the TRPV1 antagonist AMG9810 (10 µM) or the TRPA1 antagonist HC-030031 (HC; 1, 10, or 100 µM) was included in the extracellular solution for the 1.5-minute baseline recording and remained present during subsequent cephalexin exposure.

### Molecular Docking of β-Lactam Antibiotics to the TRPA1

Molecular docking was performed to compare the predicted interactions of cephalexin, amoxicillin, and JT010 with the TRPA1 binding pocket surrounding Cys621. JT010, a potent TRPA1 agonist known to activate the channel through covalent modification of Cys621, was included as a reference compound for evaluating the position of cephalexin and amoxicillin. Ligands were constructed in ChemDraw 22.0.0.22 (Revvity Signals Software). Ligands were modeled as pre-reacted covalent adducts to represent their structure after binding to the cysteine side chain. Energy minimization was done using Avogadro 1.2.0 *(46)* and AutoDock Tools 1.5.7 was used to prepare the ligands and receptor *(47)*. Docking was completed using Autodock Vina *(48, 49)*, with the Cys621 side chain treated as flexible to allow sampling of alternative conformations within the binding pocket. The holo TRPA1 structure (PDB: 6PQO) with CYS621 removed was used as the receptor for docking simulations. Docking poses were visualized using PYMOL 3.1 *(50)* in the same holo TRPA1 structure with CYS621 restored. Poses were evaluated based on AutoDock Vina docking scores and positioning of each ligand’s reactive carbon relative to Cys621. The positioning of cephalexin and amoxicillin was compared with that of JT010 as a reference ligand.

### Animal behavior testing

#### Animals

C57BL/6 mice were randomly administered vehicle or one of four different antibiotics via oral gavage. Antibiotic doses were selected based on clinically relevant dosing and/or previously reported *in vivo* exposures *(51–57)*. The doses and number of animals used for each behavioral assay are described below.

#### Facial grimace assessment

Ongoing pain behavior was assessed in 45 mice (25 male and 20 female) using the Mouse Grimace Scale *(58)*. Mice received vehicle or one of the two doses of each antibiotic: amoxicillin (30 or 90 mg/kg), azithromycin (3.5 or 10.5 mg/kg), cephalexin (15 or 45 mg/kg), or doxycycline (2 or 6 mg/kg). An individual blinded to treatment group assessed mice at baseline and at 30, 60, 90, and 120 min following antibiotic administration. Orbital tightening, nose and cheek bulge, ear position and whisker changes were scored from 0-2 and a single average was calculated for each timepoint per mouse.

#### Abdominal sensitivity assessment

Abdominal mechanical sensitivity *(59)* was assessed using a single von Frey filament (0.07 g) in 27 male mice. For this assay, only the lower dose of each antibiotic was tested: amoxicillin (30 mg/kg), azithromycin (3.5 mg/kg), cephalexin (15 mg/kg), or doxycycline (2 mg/kg). The filament was applied to the abdominal region five times at each testing time point, and the number of positive responses was recorded. Abdominal sensitivity was assessed at baseline and 30 and 120 min following antibiotic administration. Three mice exhibited an 80% response rate at baseline and were excluded from subsequent analysis.

### Fecal water content

Fecal sample water content was assessed 120-180 minutes following antibiotic administration in the abdominal sensitivity testing cohort. Fresh fecal pellets (2-3 per mouse) were collected and immediately weighed to obtain wet weight. Samples were then dried at 50 °C for 24 hours and subsequently weighed to obtain dry weight. Fecal water content was calculated as percentage of fecal wet weight attributable to water: [(wet weight - dry weight)/wet weight] x 100. Percent change in fecal water content from pre-to post-treatment was calculated for each animal. Animals were excluded if feces were not collected before the end of the 180 minute window.

### Mitochondrial superoxide (MitoSOX) imaging

Mitochondrial reactive oxygen species (ROS) production was assessed using the mitochondrial superoxide indicator MitoSOX Red. Mouse DRG neurons were cultured as described above and used within 24 hours of plating. Neurons were incubated with MitoSOX Red (5 µM final concentration) in extracellular buffer for 10 minutes at 37°C protected from light. Following dye loading, cells were washed and exposed to doxycycline (5 µg/mL) in for 5 minutes. For antioxidant experiments, cells were treated with NAC (10 mM) during the 5-minute doxycycline exposure. Vehicle-treated cells underwent the same experimental protocol in the absence of doxycycline. Following treatment, cells were imaged using the Nikon Eclipse Ti200 inverted microscope. Regions of interest were drawn around neuronal somata and mean fluorescence intensity was quantified using NIS-Elements. The threshold for MitoSOX responsiveness was defined as the 95^th^ percentile of fluorescence intensity observed in the vehicle-treated neurons. Neurons with fluorescence intensity above this threshold were classified as MitoSOX responders. The percentage of MitoSOX-responsive neurons was calculated for each treatment condition. MitoSOX fluorescence intensity was also analyzed across the total population of neurons, regardless of responder classification.

### Quantification and statistical analysis

For calcium imaging experiments, responses were quantified as the proportion of responsive neurons under each condition, with responsive cells defined based on predefined threshold of change (>25%) in the FURA-2 340/380 ratio. Comparison of the proportions between groups were performed using two-tailed Fisher’s exact tests. For continuous measures (e.g., peak 340/380 and area under the curve), data are presented as mean±SEM and were analyzed using one-way ANOVAs with appropriate multiple comparisons tests when data were normally distributed, or Kruskal–Wallis tests when normality was not met, as specified in the figure legends. Behavioral data were analyzed using one-way ANOVA with Dunnett’s multiple-comparisons tests or one-sample *t* tests, as appropriate. Pre- and post-treatment fecal water content was compared using paired *t* tests, and percent change across treatment groups was analyzed using Welch’s one-way ANOVA. MitoSOX responder proportions were compared using Fisher’s exact tests, while fluorescence intensity was analyzed using one-way ANOVA with Tukey’s multiple-comparisons test. All statistical analyses were performed using GraphPad Prism (version 10). Statistical significance was defined as p < 0.05. Significance levels are indicated as follows: * p < 0.05, ** p < 0.01, *** p < 0.001, **** p < 0.0001.

## List of Supplementary Materials

Fig. S1 to S4

Table S1 to S2

## Acknowledgments

The authors thank the organ donors and their families for their generous gift and Southwest Transplant Alliance for supporting tissue recovery work.

## Author contributions

Conceptualization: ANP, KES

Methodology: ANP, JBL, JMB, KJT, KES

Investigation: ANP, JBL, JMB, LMC, MAS, MLP, KTW, KES

Visualization: ANP, KTW, KJT, KES

Funding acquisition: KES, TJP

Writing – original draft: ANP

Writing – reviewing and editing: JBL, JMB, LMC, MAS, MLP, KTW, KJT, TJP, KES

## Data and materials availability

All data are available in the main text or supplementary materials.

