## Supplementary material for "Antibiotics modulate activity of mouse and human dorsal root ganglia neurons": Fig S1, S2, S3, S4 and Tables S1, S2

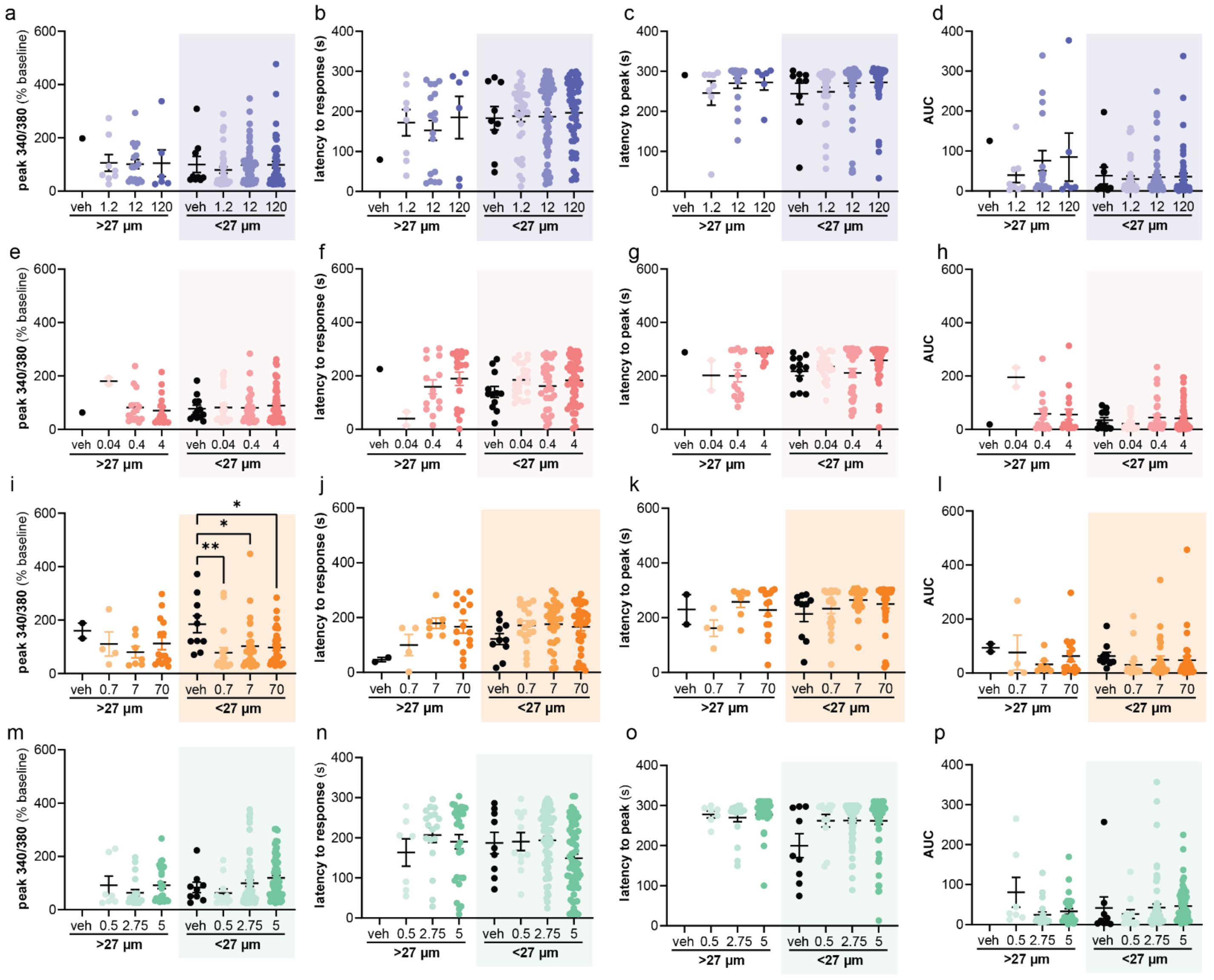

**Figure S1. Antibiotics do not change magnitude of response or response kinetics in mDRG neurons *in vitro*.** Comparison of antibiotic response peak magnitude, response latency to drug, peak response time, and area under the curve of **a.-d.**, cephalexin (1.2, 12, 120 μg/mL), **e.-h.**, azithromycin (0.04, 0.4, 4 μg/mL), **i.-l.**, amoxicillin (0.7, 7, 70 μg/mL), or **m.-p.**, doxycycline (0.5, 2.75, 5 μg/mL). Large (>27 μm) and small diameter soma (≤27 μm) were analyzed independently. Numbers on top of or within bars indicate responsive neurons/total neurons analyzed; n=4 mice per drug. Statistical comparisons versus vehicle were performed using two-tailed Fisher's exact tests within each size group; **P*< 0.05, ***P* < 0.01, ****P* < 0.001, **** *P*< 0.0001.

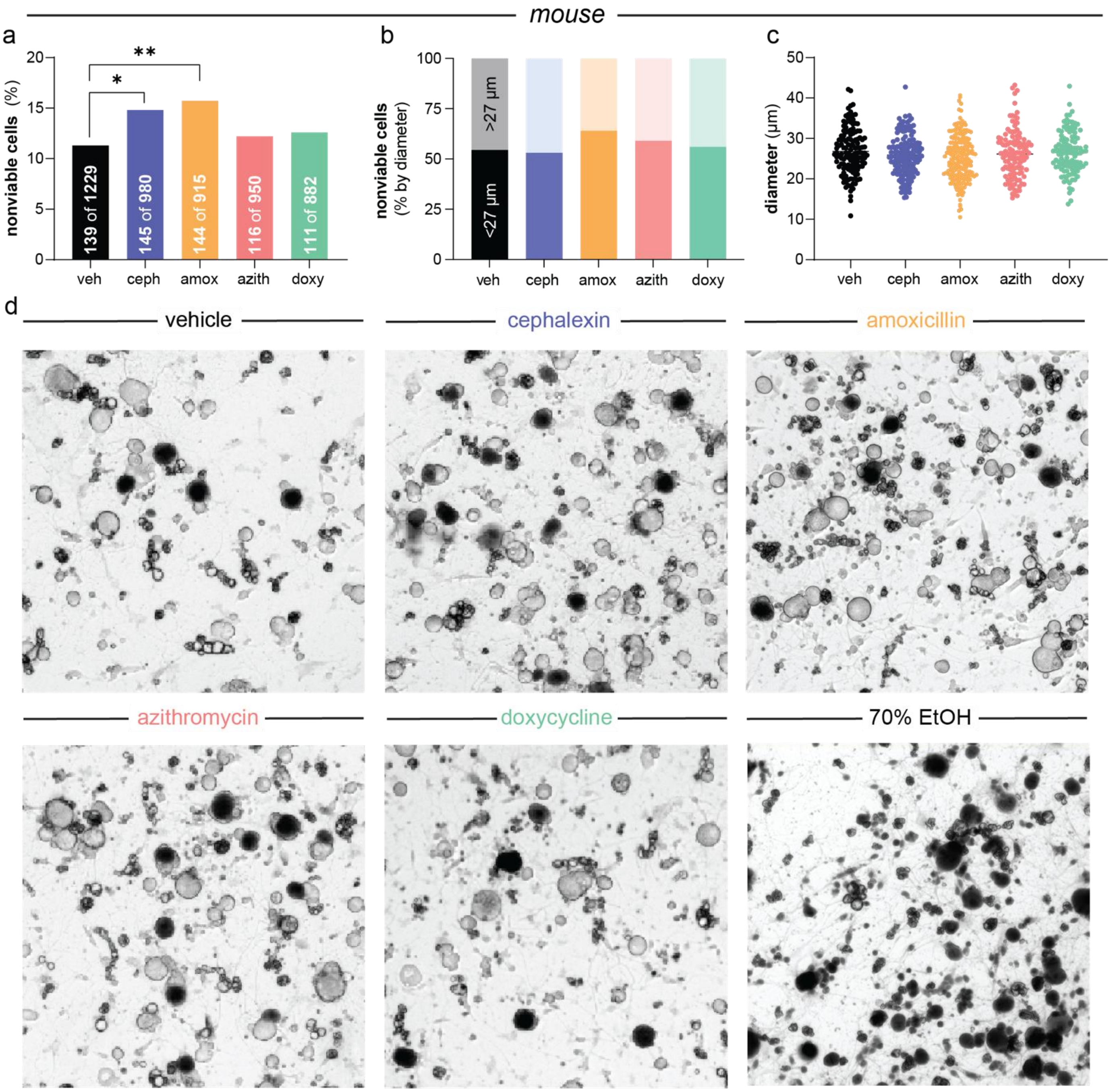


**Figure S2. Cephalexin and amoxicillin induce modest cell death in mouse DRG neurons *in vitro*. a.**, Quantification of nonviable cells after 5-minute exposure to antibiotics or control. **b.**, Percentage of nonviable cells separated large (>27 µm) and small diameter soma (≤27 µm). cephalexin (12 µg/mL), **c**., diameter of all the cells across antibiotic drugs. **d.**, representative images of mDRG neurons exposed to vehicle, cephalexin (12 µg/mL), amoxicillin (7 µg/mL), azithromycin (0.4 µg/mL), doxycycline, and 70% ethanol (positive control). azithromycin or **e.**, doxycycline (5 µg/mL). Statistical comparisons versus vehicle were performed using two-tailed Fisher's exact tests; *$P$< 0.05, **$P$ < 0.01, ***$P$ < 0.001, **** $P$< 0.0001.

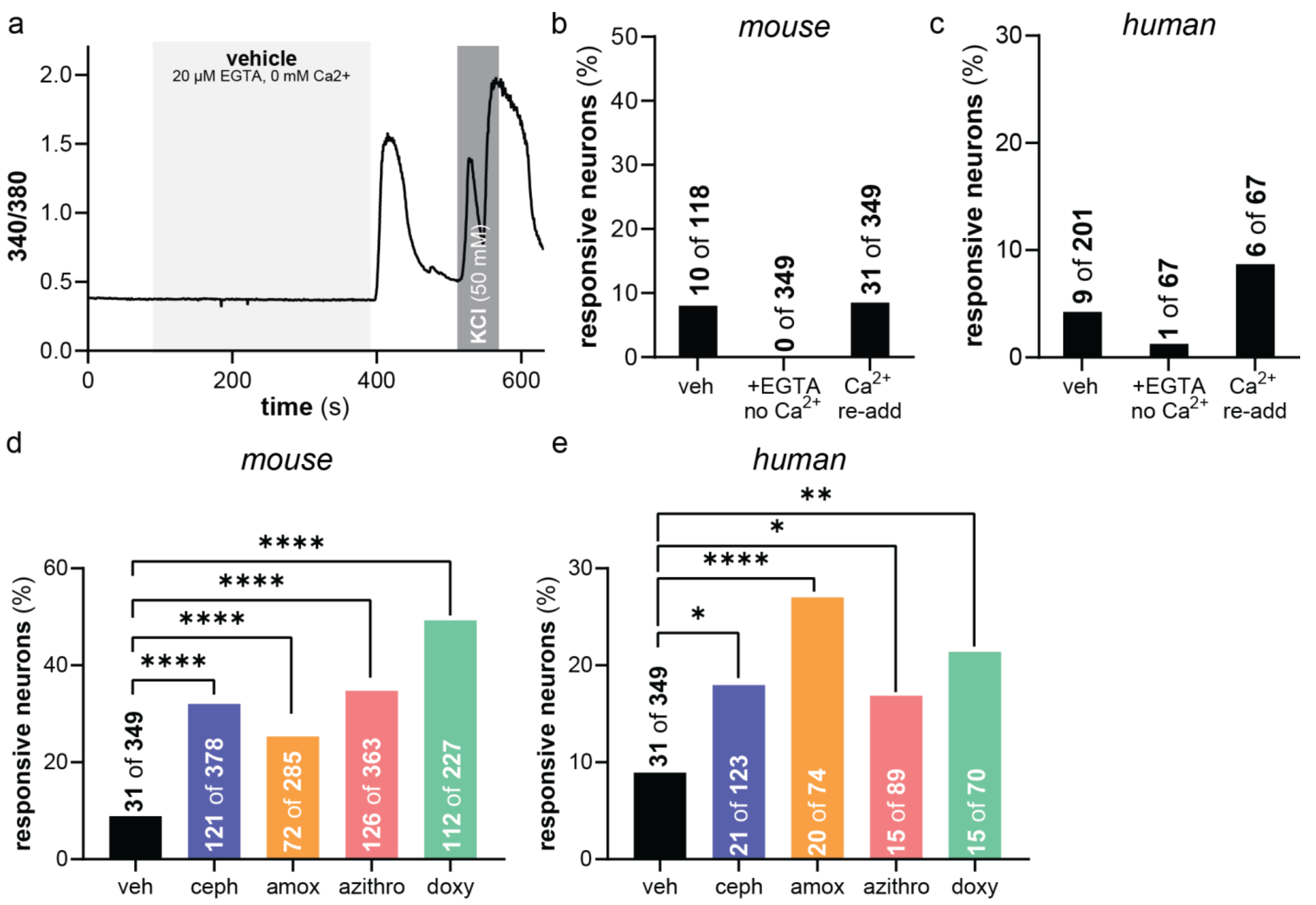


**Figure S3. Re-addition of calcium to vehicle exposed mDRG and hDRG neurons does not produce robust changes in the number of responsive neurons. a.**, Representative calcium transients from mDRG neurons exposed to vehicle. Quantification of **b.**, mDRG and **c.,** hDRG neurons that responded during each phase of imaging (veh alone, +EGTA/0 $Ca^{2+}$, and $Ca^{2+}$ re-addition). Quantification of **d.,** mDRG and **e.,** hDRG neurons that respond to antibiotics during Ca2+ re-addition compared to vehicle. Statistical comparisons were performed using two-tailed Fisher's exact tests; *$P$<0.05, **$P$<0.01, ***$P$<0.001, ****$P$<0.0001.

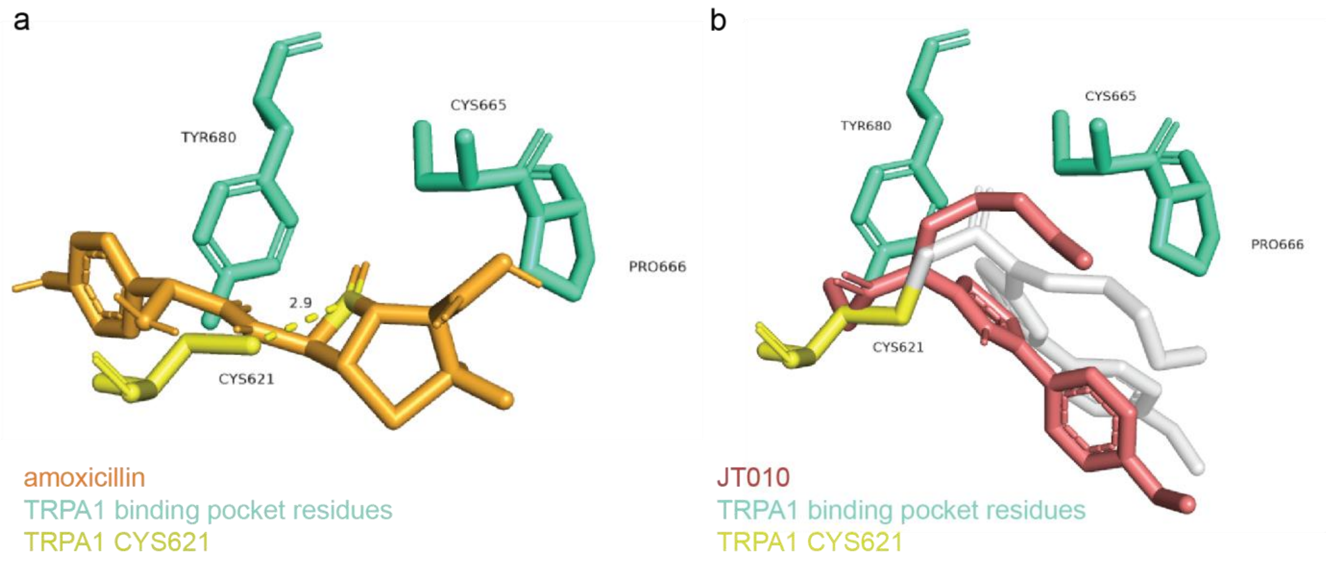


**Figure S4. Comparative docking of JT010 and amoxicillin with TRPA1.** Predicted docking poses of **a.,** the established TRPA1 agonist JT010 and **b.,** amoxicillin within TRPA1 (PDB: 6PQO), with Cys621 modeled as a flexible residue. Distances between each ligand and Cys621 are indicated.

| UTD ID | Age | Sex | COD | DRG Used | DIV | Experiment |
|---|---|---|---|---|---|---|
| UTD-DN0405 | 19 | M | Anoxia/Drug Intoxication | T12, L4 (2X) | 3 | Amox, Ceph, ENH |
| UTD-DN0410 | 35 | M | Anoxia/Drug Intoxication | T11, T12, L5 | 4 | Azithro, Dox, ENH |
| UTD-DN0423 | 35 | M | Head Trauma/Blunt Injury/MVA | L3 (2X), L4 (2X) | 3 | Amox, Dox, ENH |
| UTD-DN0426 | 33 | F | Anoxia/Asphyxiation | L5 (2X) | 6 | Azithro, Ceph, ENH |
| UTD-DN0429 | 48 | M | CVA/Stroke | L4, L5 | 4 | All |
| UTD-DN0464 | 33 | M | Anoxia/Asphyxiation/Suicide | L3 (2X) | 3 | Amox, Azithro, Dox, Ceph (EGTA) |
| UTD-DN0466 | 27 | M | Head Trauma/Blunt Injury/MVA | T12, L4 | 6 | Amox, Azithro, Dox, Ceph (EGTA) |
| UTD-DN0473 | 33 | F | CVA/Stroke | L3 (2X) | 4 | Amox, Azithro, Dox, Ceph, ENH (EGTA) |
| UTD-DN0493 | 54 | M | CVA/Stroke | L1, L4 | 3 | Dox, Ceph, Dox + RR, Ceph + RR |
| UTD-DN0495 | 26 | F | Head Trauma | T7, T11, T12, L3 | 5 | Amox, Dox, Dox + RR, Amox + RR |
| UTD-DN0498 | 48 | M | Head Trauma | L1 | 5 | Azithro, Azithro + RR |
| UTD-DN0515 | 48 | F | Anoxia | L4 (2X) | 4 | Azithro + RR |
| UTD-DN0522 | 42 | F | Cardiac Arrest | T11, L2, L3, L4 | 3 | Amox, Amox + RR |
| UTD-DN0527 | 46 | M | Anoxia | T12, L2 (2X), L3, L4 | 5 | ENH + EGTA, Azithro + RR, Ceph + RR |

**Table S1. Organ donor demographics and experimental details for human dorsal root ganglion calcium imaging.** Donor information includes age, sex, cause of death (COD), dorsal root ganglion (DRG) level, days in vitro (DIV) at the time of calcium imaging, and experimental conditions tested. M, male; F, female; Amox, amoxicillin; Ceph, cephalexin; Dox, doxycycline; Azithro, azithromycin; ENH, bath solution; RR, ruthenium red.

| Ligand | Docking Score |
| --- | --- |
| **Cephalexin** | -7.024 |
| **Amoxicillin** | -7.582 |
| **JT010** | -5.872 |

**Table S2. Docking analysis of cephalexin, amoxicillin, and JT010 with TRPA1.** Docking was performed using the TRPA1 structure (PDB: 6PQO) with Cys621 modeled as a flexible residue. Docking scores and predicted distances between each ligand and Cys621 are shown. More negative docking scores indicate more favorable predicted binding. JT010, a known covalent ligand of Cys621, was included as a reference.
